# Learning Single-Cell Perturbation Responses using Neural Optimal Transport

**DOI:** 10.1101/2021.12.15.472775

**Authors:** Charlotte Bunne, Stefan G. Stark, Gabriele Gut, Jacobo Sarabia del Castillo, Kjong-Van Lehmann, Lucas Pelkmans, Andreas Krause, Gunnar Rätsch

## Abstract

The ability to understand and predict molecular responses towards external perturbations is a core question in molecular biology. Technological advancements in the recent past have enabled the generation of high-resolution single-cell data, making it possible to profile individual cells under different experimentally controlled perturbations. However, cells are typically destroyed during measurement, resulting in *unpaired* distributions over either perturbed or non-perturbed cells. Leveraging the theory of optimal transport and the recent advents of convex neural architectures, we learn a coupling describing the response of cell populations upon perturbation, enabling us to predict state trajectories on a single-cell level. We apply our approach, CellOT, to predict treatment responses of 21,650 cells subject to four different drug perturbations. CellOT outperforms current state-of-the-art methods both qualitatively and quantitatively, accurately capturing cellular behavior shifts across all different drugs.

## 1 Introduction

Characterizing and modeling perturbation responses at the single-cell level from non-time resolved data remains one of the grand challenges of biology. It finds applications in predicting cellular reactions to environmental stress or a patient’s response to drug treatments. Accurate inference of perturbation responses at the single-cell level allows us, for instance, to understand how and why individual tumor cells evade cancer therapies (Frangieh et al., 2021). More generally, it deepens the mechanistic understanding of the molecular machinery determining the respective responses to perturbations.

Cell responses to perturbations such as drugs are highly heterogeneous in nature (Liberali et al., 2014), determined by many factors, including the preexisting variability in the abundance and localization of molecular entities, such as RNA or proteins (Shaffer et al., 2017), cellular states (Kramer and Pelkmans, 2019), or the cellular microenvironment (Snijder et al., 2009). To effectively predict the drug response of a patient during treatment, it is thus crucial to incorporate the *molecular subpopulation structure* of the cell populations into the analysis.

A key difficulty in learning perturbation responses is that a cell (usually) must be destroyed to measure its state, meaning that it is only possible to measure a cell state either before or after a perturbation is applied. The typical experimental setup divides a set of cells into subsets to which individual perturbations are applied. Hereby, a subset of cells remains unperturbed, allowing us to measure the base state of the population. So while we do not have access to a set of paired control/perturbed single-cell observations, we do have access to samples of distributions of control/perturbed cell states.

Previous methods to approximate single-cell perturbation responses fall short of solving this highly complex *pairing problem* while, at the same time, accounting for cellular heterogeneity and the strong subpopulation structure of cell samples. Despite incorporating cell heterogeneity, mechanistic models do not recover cellular response trajectories, instead of predicting factors such as cell viability or response variables in the data in order to predict drug efficacy (Snijder et al., 2012; Berchtold et al., 2018; Green and Pelkmans, 2016). Linear models (Dixit et al., 2016), on the other hand, are unable to capture complex and inhomogeneous population responses upon perturbation. Current state-of-the-art methods (Lopez et al., 2018; Lotfollahi et al., 2019; Yang et al., 2020) predict perturbation responses via linear shifts in a learned low-dimensional latent space. While capturing nonlinear cell-type-specific responses, their use of linear interpolations cause them to resolve the alignment problem with the challenging task of learning representations that are invariant to their perturbation status. A similar matching problem was considered in Stark et al. (2020) for matching cell populations that are profiled with different profiling technologies.

This work proposes CellOT, a novel approach to predict single-cell perturbation responses by uncovering couplings between control and perturbed cell states while accounting for heterogeneous subpopulation structures of molecular environments. We achieve this by utilizing the theory of optimal transport, which provides natural geometry and mathematical tools to manipulate probability distributions. To this end, we learn a robust optimal transport map describing how the distribution of control cells connects to the distribution of perturbed cells. Utilizing recent developments of neural optimal transport (Makkuva et al., 2020), we learn a general optimal transport coupling for each perturbation, allowing us to predict behavioral changes of incoming single-cell samples, e.g., of another patient, using parameterizations learned for the previous cohort. We demonstrate CellOT’s effectiveness by deploying it to learning cellular responses to different cancer drugs and tumor combination therapies. An overview of our approach is illustrated in Figure 1.

**Figure 1:**
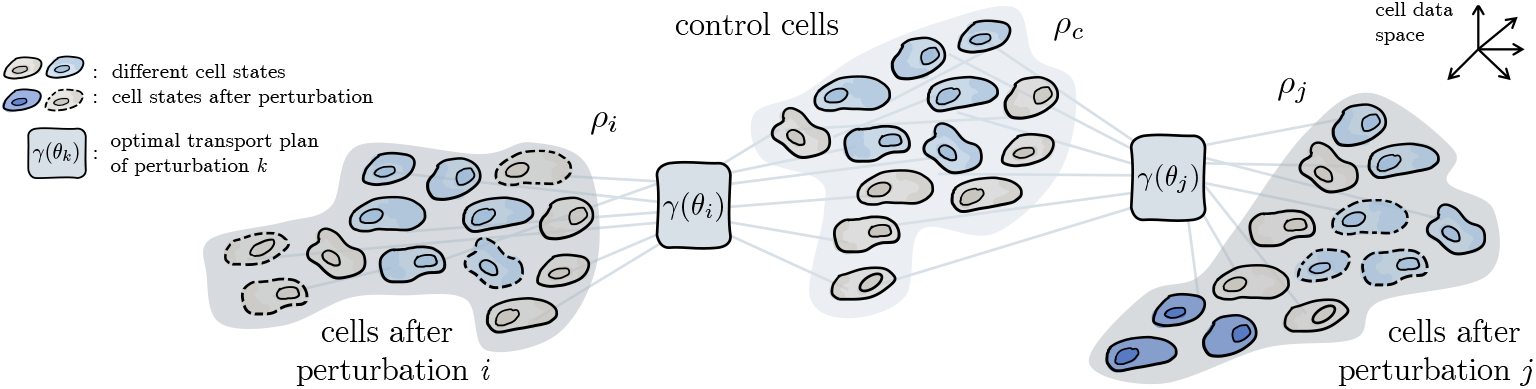
Learning single-cell perturbation responses. We aim to recover a mapping from control cell distributions *ρ_c_* to some perturbed cell distribution *ρ_i_* or *ρ_j_* by learning the corresponding neural optimal transport map *γ*(*θ_k_*), parameterized by *θ_k_*, from the observed distribution of untreated cells and the set of cells observed after the perturbation is applied.

Optimal transport has previously been applied in the domain of single-cell biology to uncover trajectories of single-cell reprogramming and to link rich, non-spatially-resolved with sparse, spatially resolved measurements (Schiebinger et al., 2019; Cang and Nie, 2020; Demetci et al., 2020; Huizing et al., 2021; Lavenant et al., 2021; Zhang et al., 2021). Here we apply optimal transport to a new data modality, consisting of cell morphological measurements and multiplexed protein state measurements obtained by 4i (Gut et al., 2018) from large populations of cancer cells exposed *in vitro* to different drugs used in the clinic.

## 2 Background

### 2.1 Optimal Transport

Optimal transport plays dual roles as it induces a mathematically well-characterized distance measure between distributions besides providing a geometry-based approach to realizing couplings between two probability distributions. Let 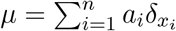 and 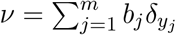 be two discrete probability measures in 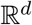. The optimal transport (OT) problem (Kantorovich, 1942) reads

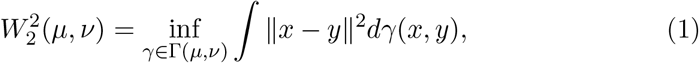

where the polytope Γ(*a*, *b*) is 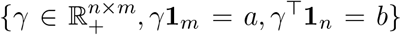 describes the set of all couplings *γ* between *μ* and *v*. The optimal transport plan *γ* thus corresponds to the coupling between two probability distributions minimizing the overall transportation cost. Computing optimal transport distances in (1) involves solving a linear program, and thus their computational cost is prohibitive for large-scale machine learning problems. Regularizing objective (1) with an entropy term results in significantly more efficient optimization (Cuturi, 2013),

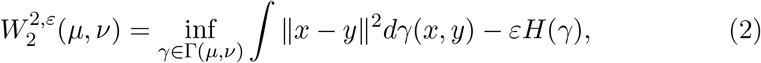

with entropy 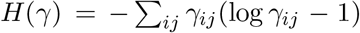 and parameter *ε* controlling the strength of the regularization. 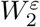 is further differentiable w.r.t. its inputs and thus serves as a loss function in machine learning applications.

Problem (1) denotes the primal formulation for the Wasserstein-2 distance. The corresponding dual introduced by Kantorovich in 1942 is a constrained concave maximization problem defined as

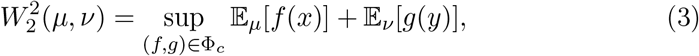

where the set of admissible potentials is 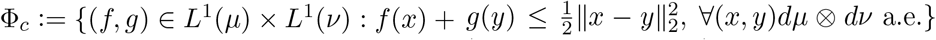 (Villani, 2003, Theorem 1.3). Villani (2003, Theorem 2.9) further simplifies the dual problem (3) over the pair of functions (*f, g*) to

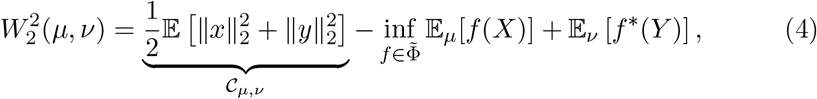

where 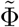 is the set of all convex functions in *L*^1^ (*dμ*) × *L*^1^(*dv*), *L*^1^(*μ*) ≔ {*f* is measurable & ∫ *fdμ* < ∞}, and *f*\*(*y*) = sup_*x*_〈*x, y*〉 – *f*(*x*) is *f*’s convex conjugate. Villani (2003, Theorem 2.9) then proves the existence of an optimal pair (*f*, *f*\*) of lower semi-continuous proper conjugate convex functions on 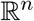 minimizing (3).

### 2.2 Convex Neural Networks

In order to parameterize convex spaces such as 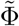 in (4), we need neural networks which are convex w.r.t. to their inputs. One example are input convex neural networks (ICNN) introduced by Amos et al. (2017). ICNNs are based on fully-connected feed-forward networks that ensure convexity by placing constraints on their parameters. An ICNN with parameters 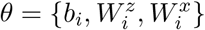 represents a convex function *f*(*x*; *θ*) and, for a layer *i* = 0 … *L* – 1, is defined as

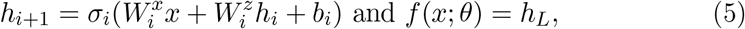

where activation functions *σ_i_* are convex and non-decreasing, and elements of all 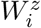 are constrained to be nonnegative. Despite their constraints, ICNNs are able to parameterize a rich class of convex functions. In particular, Chen et al. (2019) provide a theoretical analysis that any convex function over a convex domain can be approximated in sup norm by an ICNN. Huang et al. (2021) further extend ICNNs from fully-connected feed-forward neural networks to convolutional neural architectures.

### 2.3 Neural Optimal Transport

Despite existing numerical approximations of the optimal transport distance and the corresponding optimal coupling (Cuturi, 2013; Aude et al., 2016) (2), recent efforts have investigated neural network-based approaches as fast and scalable approximations to (1). Taghvaei and Jalali (2019) consider solving (4) by parameterizing *f* with an ICNN and solve for *f*\* at each step, which has a high computational cost. Makkuva et al. (2020) extend this work by approximating *f*\* with another ICNN *g*, which scales well but transforms the problem into a min-max optimization task. Huang et al. (2021) introduce a novel, OT-inspired parameterization of normalizing flows utilizing ICNNs. Korotin et al. (2021) provide a detailed comparison of the current state of neural optimal transport solvers. Furthermore, convex neural architectures have been utilized to parameterize Wasserstein gradient flows (Bunne et al., 2021; Alvarez-Melis et al., 2021; Mokrov et al., 2021).

## 3 Model

Recent high-throughput methods provide great insights on how cell populations respond to various perturbations on the level of individual cells. The provided data, however, is non-time-resolved and unaligned. Hence, snapshots taken of biological samples before and after perturbations do not provide information on single-cell trajectories. Perturbations might include the application of drugs affecting molecular functions in cells, or changes in the cellular environment causing shifts in biological signaling, thus impacting cells and their states in various ways. In the following, we describe our approach, which uncovers single-cell perturbation responses by predicting couplings between control and perturbed cell states. Hereby, let 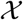 denote the biological data space spanned by cell morphology and gene expression features. We then treat a cell’s response to perturbation *k* as an evolution in a high-dimensional space of cell states 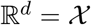.

### 3.1 Recovering Perturbation Effects via Optimal Transport

Given a dataset of *n* observations 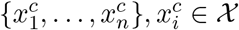 drawn from 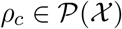, the distribution of cells before applying a perturbation, we aim to learn the distribution of cells 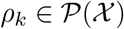 upon some perturbation *k*, given a set of separate samples 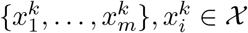.

Perturbation responses of cells are dynamic: after applying perturbation *k*, cell states evolve over time and thus can be modeled as a stochastic process on the cell data space. Despite this time-resolved nature of single-cell responses, we only have access to the distributions of cell states before, *ρ_c_*, and after injecting perturbation *k*, *ρ_k_*. We thus aim at understanding the underlying stochastic process without access to time-resolved perturbation responses by uncovering the coupling *γ* between *ρ_c_* and *ρ_k_*. Given prior biological knowledge, we can assume that cells do not drastically alter their phenotype w.r.t. morphology and gene expression pattern. We thus posit that the evolution of probability distributions of single-cells upon perturbation can be modeled via the mathematical theory of optimal transport. The coupling *γ* then corresponds to an optimal transport plan (1) between *ρ_c_* and *ρ_k_*.

Following Makkuva et al. (2020), we infer the optimal coupling *γ* (1) between *ρ_c_* and *ρ_t_*. Thus, instead of computing a coupling individually for each pair of cell samples using existing solvers (Cuturi, 2013), we learn a parameterized optimal transport map using neural networks. The parameterized OT coupling then serves as a robust predictor for cellular distribution shifts upon perturbations on unseen samples 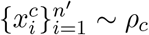, i.e., of another patient.

### 3.2 Parametrization of the Optimal Transport Coupling

Directly learning the optimal transport map in the primal (1) and dual (3) is notoriously difficult. Instead, Makkuva et al. build upon celebrated results by Knott and Smith (1984) and Brenier (1991), which relate the optimal solutions for the dual form (3) and the primal form (1), to derive a min-max formulation replacing the convex conjugate in (4) (Makkuva et al., 2020, Theorem 3.3)

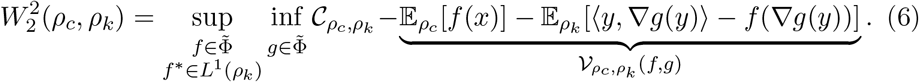

We can further relax the constraint 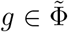 to *L*^1^(*ρ_k_*), as a function *g* ∈ *L*^1^(*ρ_k_*) minimizing (6) is convex and equal to *f* * for any convex function *f*. In order to learn the resulting optimal transport, i.e., the solution of the minimization problem in (6), Makkuva et al. (2020) parametrize both dual variables *f* and *g* using input convex neural networks (§ 2.3) (Amos et al., 2017). The resulting approximate Wasserstein distance is thus defined as

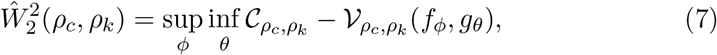

where *θ* and *ϕ* are the parameters of each ICNN. The resulting 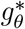 produces an approximate optimal transport plan 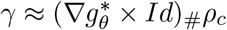.

### 3.3 Predicting Perturbation Effects via CellOT

The framework described above allows us to recover couplings between control 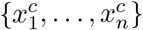 and perturbed cells 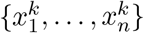, giving insights into cellular response trajectories upon application of a perturbation *k*. Given a set of perturbations *K*, and sample access to the control distribution *ρ_c_* as well as distributions *ρ_k_* for each perturbation *k* ∈ *K*, CellOT learns the optimal pair of dual potentials 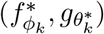 for each perturbation *k*. Given parametrizations of the convex potentials for each *k*, CellOT then predicts the transformation of a control cell 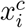 upon perturbation *k* via 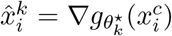, i.e., samples following the predicted perturbed distribution 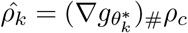. CellOT thus provides a general approach to predict state trajectories on a single-cell level, as well as understand, how heterogeneous subpopulation structures evolve under the impact of external factors.

## 4 Evaluation

We evaluate CellOT on the task of predicting single-cell drug responses for drugs with different molecular effects, using melanoma cell lines profiled by the 4i technology (Gut et al., 2018).

### 4.1 Datasets

4i is an imaging technology that detects protein abundance by attaching a fluorescent tag designed to bind to a target protein and then measuring the fluorescence intensity of this tag. An iterative staining and washing procedure allows for the capture of multiple tags. Additionally, an image processing pipeline extracts morphological features, such as cell perimeter and area and detects the cell nucleus. We considered four common cancer therapies for this works since they target different biological processes. Erlotinib is an inhibitor of the epidermal growth factor receptor (EGFR) tyrosine kinase, Imatinib inhibits the Bcr-Abl tyrosine kinase, and Trametinib is an inhibitor of mitogen-activated extracellular signal-regulated kinase 1 (MEK1) and MEK2.

We utilized a mixture of 2 melanoma tumor cell lines (ratio 1:1) in order to image a total of 21,650 cells, of which 11,526 are in the (untreated) control state, 2,364 are treated with Erlotinib, 2,650 with Imatinib, 2,683 with Trametinib, and 2,417 are treated with a combination of Trametinib and Erlotinib, and 48 features are extracted for each cell. 22 features are morphological, and the remaining 26 are mean intensities of 13 protein markers detected both inside the cell nucleus and in the cell as a whole. Finally, we perform an 80/20 train test split for each condition and evaluate model performance on its ability to make predictions on the unseen set of control cells. More details regarding dataset preparation can be found in Appendix B.1.

### 4.2 Baselines

We compare CellOT to two other baselines, both of which attempt to add perturbation effects through the manipulation of a learned latent representation: scGen (Lotfollahi et al., 2019) computes linear shifts using latent space arithmetic to remove the source condition and add the target condition, and the conditional autoencoder, cAE, which has an architecture based on batch correction technique popular in the single-cell community, first introduced by Lopez et al. (2018). Here, one-hot encodings of batch labels (treatment conditions) are concatenated to the encoder and decoder inputs, which attempt to remove and then add condition-specific effects. More details can be found in Appendix A.

### 4.3 Evaluation Metrics

Since we lack access to the ground truth set of control and treatment observations on the single-cell level, we first analyze the effectiveness of CellOT using evaluations that operate on the level of the distribution of real and predicted perturbation states. Drug signatures are computed as the difference in means between the distribution of perturbed states and control states. We then report the ℓ_2_-distance between the drug signatures (DS) computed on the true and predicted distributions (ℓ_2_(*DS*)). We additionally consider two distributional distances: kernel maximum mean discrepancy (MMD) (Gretton et al., 2012) and entropy-regularized Wasserstein distance 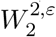 (2) (Cuturi, 2013). MMD is computed using the RBF kernel and averaging over the length scales 0.5, 0.1, 0.01, and 0.005; 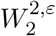 is computed with *ε* = 0.5.

### 4.4 Results

For each drug perturbation, all models predict the perturbed cell states from the set of held-out of control cells. Differences between the distribution of perturbed cells and predicted cells are shown in Table 1. CellOT significantly outperforms all baselines on all three metrics. Qualitative assessment of the marginal distributions of control, treated, and predicted cell states provides further evidence for superior performance of CellOT over other approaches (see Figure 2 for three selected features of the Imatinib condition).

**Table 1:** Performance assessment of CellOT compared to different baselines w.r.t. to Wasserstein (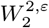, (2)) and MMD distances between the observed perturbed cells and predicted responses from control cells, as well as the predictive quality of drug signatures (see § 4.3).

| Model | Drugs |  |  |  |  |  |  |  |  |  |  |  |
| --- | --- | --- | --- | --- | --- | --- | --- | --- | --- | --- | --- | --- |
|  | Erlotinib |  |  | Imatinib |  |  | Trametinib |  |  | Trametinib and Erlotinib |  |  |
| | $\ell_2$(DS) | MMD | $W_2^{2,\varepsilon}$ | $\ell_2$(DS) | MMD | $W_2^{2,\varepsilon}$ | $\ell_2$(DS) | MMD | $W_2^{2,\varepsilon}$ | $\ell_2$(DS) | MMD | $W_2^{2,\varepsilon}$ |
| ● SCGEN | 0.41 | 0.0241 | 3.542 | 0.52 | 0.0361 | 5.811 | 0.56 | 0.0180 | 3.587 | 0.60 | 0.0163 | 3.594 |
| ● CAE | **0.05** | 0.0074 | **3.330** | 0.16 | 0.0200 | 4.512 | 0.37 | 0.0087 | 3.343 | 0.44 | 0.0122 | 3.215 |
| ● CELLOT | 0.22 | **0.0013** | 3.619 | **0.12** | **0.0010** | **3.851** | **0.14** | **0.0011** | **2.846** | **0.18** | **0.0014** | **2.796** |

**Figure 2:**
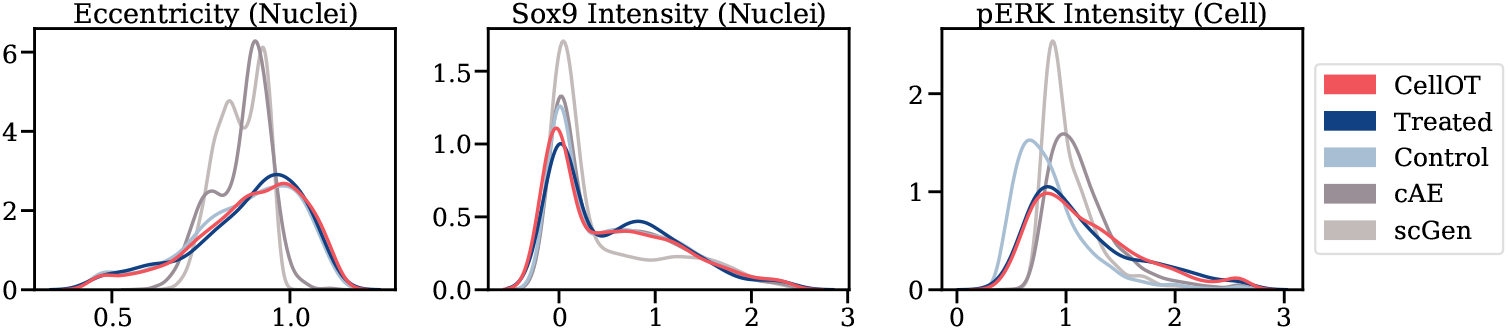
Marginal distributions of observed and predicted cell states for three selected features, i.e., a measure of the eccentricity of the nucleus, as well as Sox9 intensity level inside the nucleus and pERK intensity within the cell but outside the nucleus. The marginals of control and Imatinib distributions correspond to the observed set of untreated cells and cells treated with the Imatinib drug. The remaining distributions are calculated using the predictions of each model on the unseen set of control cells.

Next, we compared UMAP projections (McInnes et al., 2018) of the perturbed and predicted cells (see Figure 3). Predicted cells are colored by the fraction of other predicted cells in their *k* = 100 nearest neighbors. If the distribution of predicted cells matches the true distribution of perturbed cells, then we would expect the nearest neighbor of each cell to be well mixed (i.e., 0.5) across conditions. Thus, cells with values closer 1 indicate regions where the predicted distribution does not integrate with the true perturbed distribution. We conclude that predictions made by CellOT integrate well with measurements of real treated cells.

**Figure 3:**
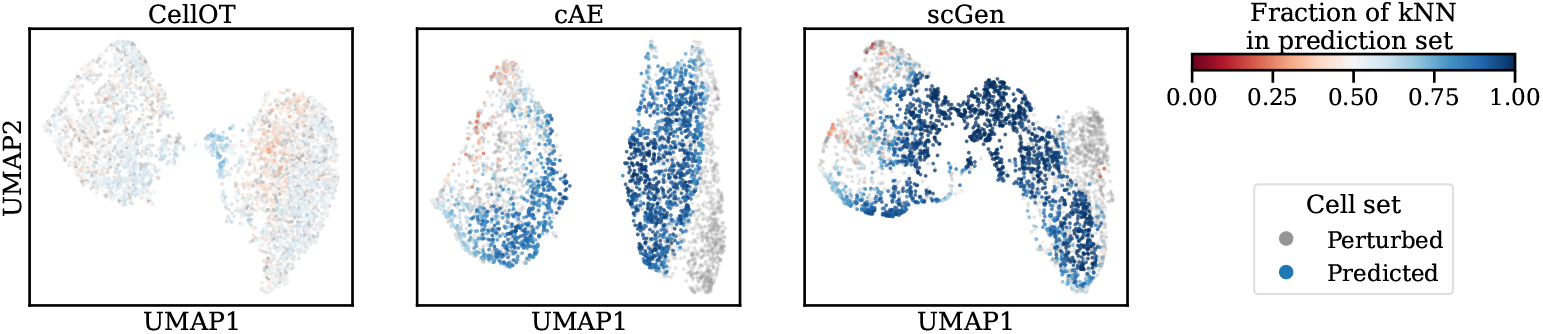
UMAP projections computed on the joint set of cells perturbed by Imatinib (grey) and predictions of each model. Model predictions are colored by the fraction of other predicted cells in their *k* = 100 nearest neighbors in data space. Predicted cells which do not share many neighbors of the true set of perturbed cells, but instead control cells take values of ≈ 1.0 (blue). Predicted cells that integrate well with true perturbed cells take values of ≈ 0.5 (white).

Finally, the previous results argue that the distribution of cells predicted by CellOT closely matches true distribution; however, they could have also been replicated by a map that assigns control cells to random treated cells. Thus, we evaluate the quality of single-cell level pairs induced from the CellOT mapping by computing the Spearman correlation of features between the control state and predicted drug state. The distribution of the correlation coefficient between the control state and the predicted state across all features of all learned maps is shown in Figure 4. The low distributional distances between CellOT-predicted cells to the true distribution of perturbed cells in conjunction with a high correlation of features with the control states paired to the predicted states demonstrate that CellOT makes sound predictions on the single-cell level, outperforming current state-of-the-art methods both qualitatively and quantitatively.

**Figure 4:**
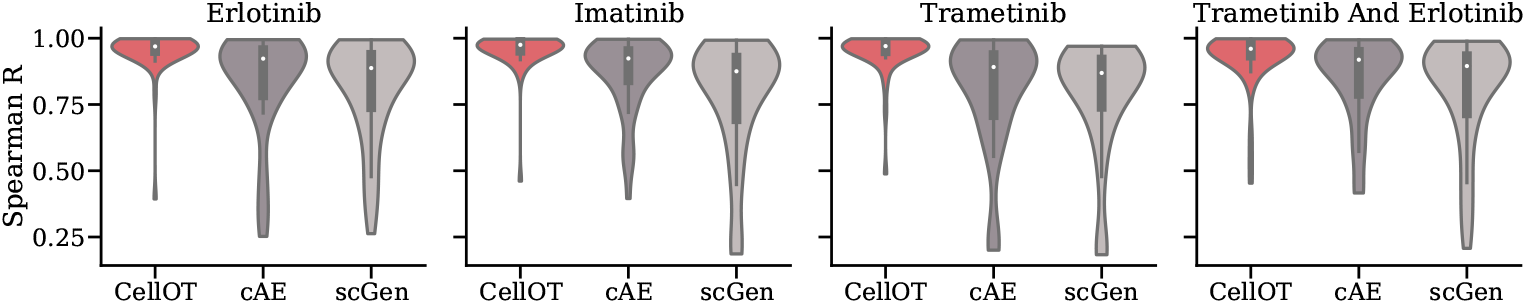
Distribution of Spearman correlation coefficients between the features of control cells and the features of its corresponding predicted state upon treatment for each considered drug Erlotinib, Imatinib, Trametinib and the combination therapy of Trametinib and Erlotinib. Low correlations imply unexpected significant differences in the feature states between prediction and control, and thus a reduced accuracy of predictive power.

**Figure 5:**
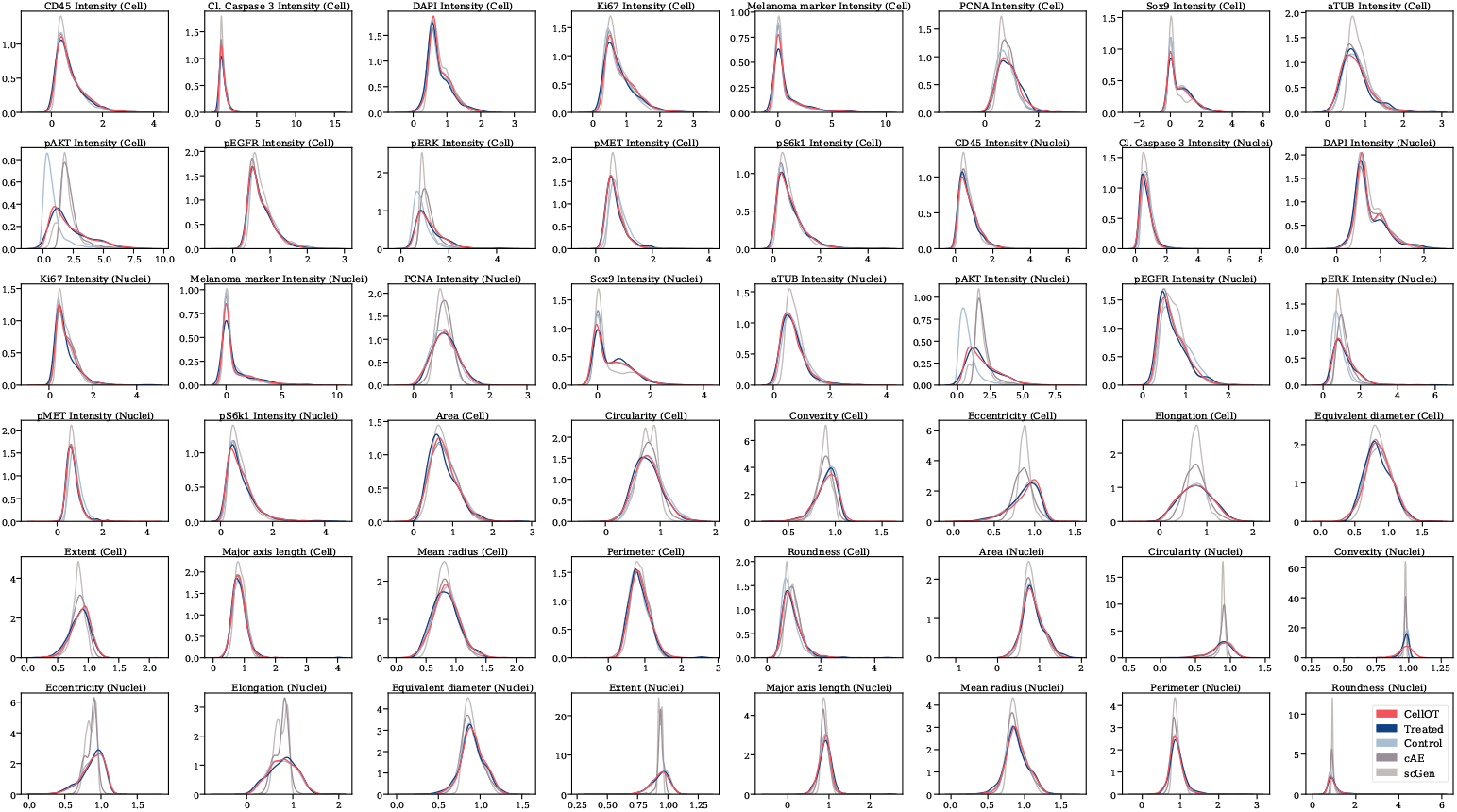
Full set of Imatinib marginals.

**Figure 6:**
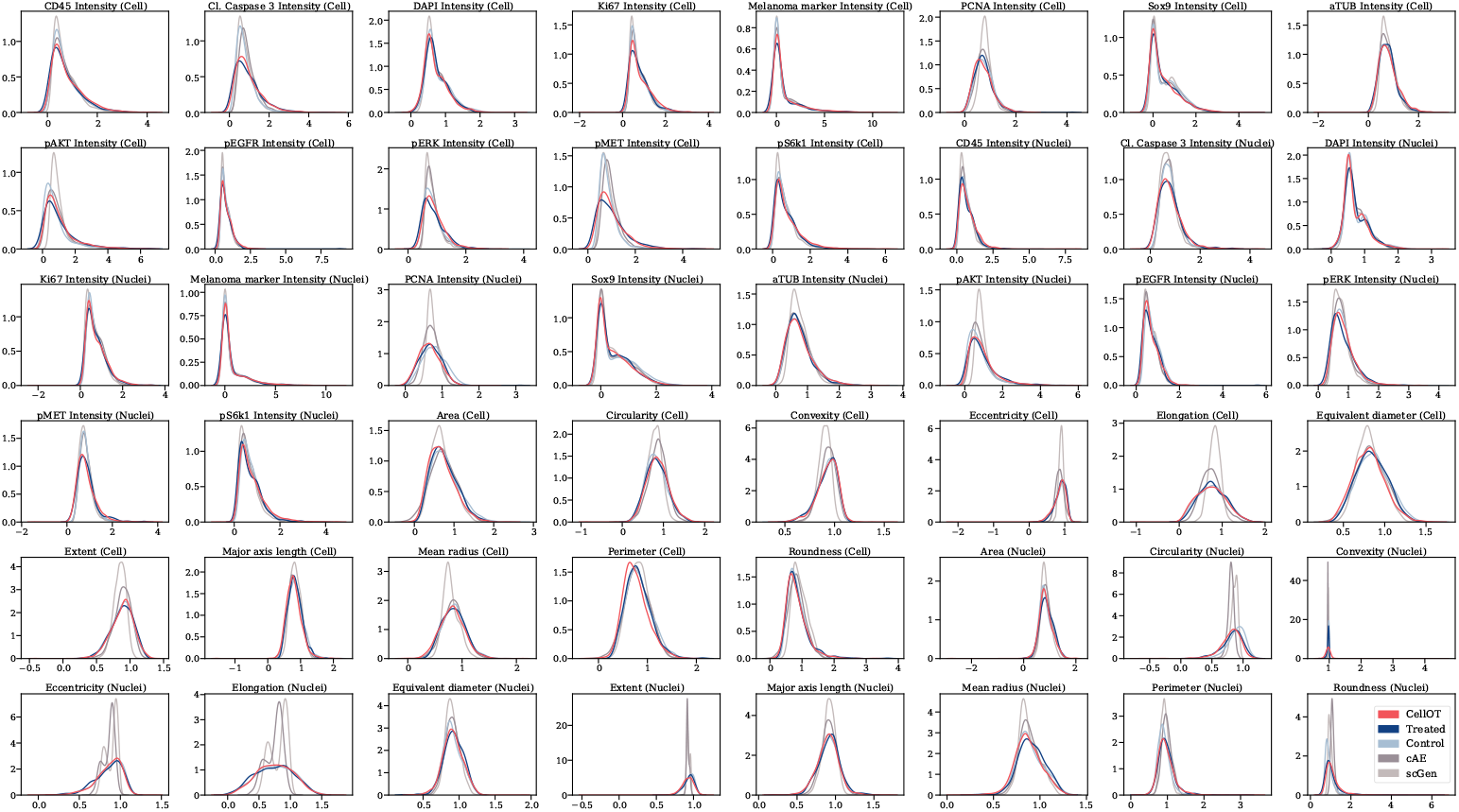
Full set of Erlotinib marginals.

## 5 Conclusion

In this paper, we present a new framework to learn single-cell perturbation responses. We approach the problem by learning an optimal transport map that is parameterized by an ICNN to push-forward the distribution of control cells onto the distribution of perturbed cells. We validate CellOT’s effectiveness through experiments on melanoma cell lines with four different drug perturbations. In the absence of ground truth, we provide various evaluation metrics to compare our method to existing approaches. While operating in the original data space, instead of relying on meaningful low-dimensional representations, CellOT performs consistently well across all perturbations, outperforming current state-of-the-art methods. The use of neural optimal transport to learn single-cell drug responses makes for an exciting avenue of future work, including its use to improve our mechanistic understanding of cell therapies, to study drug responses from patient samples, and to better account for cell-to-cell variability in large-scale drug discovery efforts.

## Acknowledgments

We are grateful to Hugo Yèche and Ximena Bonilla for their fruitful comments, corrections, and discussions. C.B. and A.K. received funding from the Swiss National Science Foundation under the National Center of Competence in Research (NCCR) Catalysis under grant agreement 51NF40 180544. L.P. is supported by the European Research Council (ERC-2019-AdG-885579), the Swiss National Science Foundation (SNSF grant 310030_192622), the Chan Zuckerberg Initiative, and the University of Zurich. G.G. received funding from the Swiss National Science Foundation and InnoSuisse as part of the BRIDGE program as well as from the University of Zurich through the BioEntrepreneur Fellowship. K.L. and S.G.S. were partially funded by ETH Zürich core funding (to G.R.) and from the Tumor Profiler Initiative (to G.R.).

## Appendix

### A Related Work

Consider a single-cell dataset of a binary perturbation. Let {*x*_1_ … *x_n_*}, 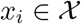, drawn from *ρ_c_* ∪ *ρ_k_* and let *c*(*i*) ∈ {0, 1} indicate the perturbation status of a single cell,

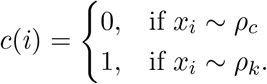

#### A.1 scGen

Given representations {*z*_1_ … *z_n_*} of {*x*_1_ … *x_n_*}, learned by an autoencoder, with encoder *ϕ* and decoder *ψ*, scGen (Lotfollahi et al., 2019) predicts a perturbation response using latent space arithmetic. Let 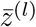 be the mean of representations in condition *l*

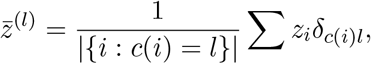

the perturbed state of *x*′ ~ *ρ_c_* is predicted as

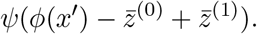

#### A.2 cAE

The conditional autoencoder is based on a batch correction technique popular within the single-cell community, first introduced by (Lopez et al., 2018). It introduces condition-specific parameters into the encoder and decoder, which attempt to remove and replace information in the data specific to their conditions. They operate by concatenating one-hot encodings of condition labels (here, perturbation status) to the inputs of the encoder and decoder. These encodings, in effect, make the bias term in the first layer of the encoder and decoder a learnable parameter specific to each condition and are thus are also considered to learn a linear shift in latent space. Given an encoder *ϕ* and decoder *ψ*, the network is trained to reconstruct cells conditioning on its true label

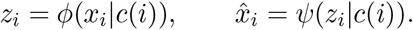

Once trained, the perturbed state of *x*′ ~ *ρ_c_* is predicted as

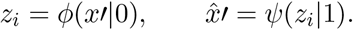

### B Dataset

#### B.1 Single-Cell Multiplex Data

Biologists have various powerful technologies at their disposal, capable of capturing multivariate single-cell measurements. High-content imaging, particularly when augmented by multiplexing abilities such as by Iterative Indirect Immunoflu-orescence Imaging (4i) (Gut et al., 2018), is ideally suited to study heterogeneous cell responses. With 4i, fluorescently labeled antibodies are iteratively hybridized, imaged, and removed from a sample to measure the abundance and localization of proteins and their modifications. Thus, 4i quickly generates large, spatially resolved phenotypic datasets rich in molecular information from thousands of treated and untreated (control) cells. Additionally to the multiplexed information 4i generates, information about cellular and nuclear morphology is routinely extracted from microscopy images (without the need for 4i) by image analysis algorithms (Carpenter et al., 2006).

Through multiplexing, 4i datasets are able to capture meaningful features related to both the treatment response heterogeneity (e.g., the phosphorylation or dephosphorylation of a kinase in a signaling pathway) and the pre-existing cell-to-cell variability (e.g., protein levels related to different cellular states or cell cycle phases) which my determine treatment response. Traditional high-content imaging datasets often need to compromise between features describing either the former or the latter and may thus struggle to provide sufficient information to pair treated and control cells accurately.

The cells were seeded in a 384-well plate, allowed to settle and adhere overnight. Drugs and Dimethyl sulfoxide as the vehicle control was added to the cells the next morning and incubated for 8 hours, after which the cells were fixed with Paraformaldehyde. Subsequently, 6 cycles of 4i were performed, for which the images were acquired with an automated high-content microscope.

All image analysis steps were performed by our in-house platform called TissueMAPS (https://github.com/TissueMAPS). The steps included illumination correction (Snijder et al., 2012), alignment of images from different acquisition cycles using Fast Fourier Transform (Guizar-Sicairos et al., 2008), segmentation of nuclei and cell outlines (Stoeger et al., 2015), as well cellular and nuclear measurements of intensity and morphology features using the scikit-image library (Van der Walt et al., 2014).

### C Experimental Details

To train all networks, we use the Adam optimizer (Kingma and Ba, 2014).

#### C.1 Baselines

To tune baseline models, we use a batch size of 128 and do a grid search over the width [16, 32] and depth [2, 3] of the encoder and decoder hidden layers, latent dimension [4, 8], dropout rate [0.0, 0.05, 0.1, 0.2] and learning rate [0.00001, 0.0001, 0.001].

For both scGen and cAE we selected a width=32, depth=2, latent dim=8, dropout=0.05. scGen uses a learning rate of 0.001, and cAE uses a learning rate of 0.0001. Both models are trained for 1024 epochs.

#### C.2 Network Architectures

As suggested by Makkuva et al. (2020), we relax the convexity constraint on *g_θ_* and instead, penalize its negative weights 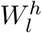

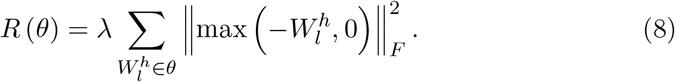

The convexity constraint on *f_ϕ_* is enforced after each update by setting negative weights of all 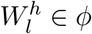 to zero. Thus the full objective then states

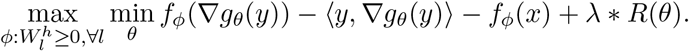

#### C.3 Hyperparameters

To learn the optimal transport maps, we use a batch size of 256, an ICNN architecture of 4 hidden layers of width 64, a learning rate of 0.0001 (*β*_1_ = 0.5, *β*_2_ = 0.9) and λ=1. The inner loop minimizing *g* runs for 10 updates to every update of *f*.

## References

D. Alvarez-Melis, Y. Schiff, and Y. Mroueh. Optimizing Functionals on the Space of Probabilities with Input Convex Neural Networks. arXiv Preprint, 2021.

B. Amos, L. Xu, and J. Z. Kolter. Input Convex Neural Networks. In International Conference on Machine Learning (ICML), volume 34, 2017.

G. Aude, M. Cuturi, G. Peyré, and F. Bach. Stochastic Optimization for Large-Scale Optimal Transport. In Advances in Neural Information Processing Systems (NeurIPS), 2016.

D. Berchtold, N. Battich, and L. Pelkmans. A systems-level study reveals regulators of membrane-less organelles in human cells. Molecular cell, 72(6): 1035–1049, 2018.

Y. Brenier. Polar Factorization and Monotone Rearrangement of Vector-Valued Functions. Communications on pure and applied mathematics, 44(4):375–417, 1991.

C. Bunne, L. Meng-Papaxanthos, A. Krause, and M. Cuturi. JKOnet: Proximal Optimal Transport Modeling of Population Dynamics. arXiv Preprint, 2021.

Z. Cang and Q. Nie. Inferring spatial and signaling relationships between cells from single cell transcriptomic data. Nature Communications, 11(1), 2020.

A. E. Carpenter, T. R. Jones, M. R. Lamprecht, C. Clarke, I. H. Kang, O. Friman, D. A. Guertin, J. H. Chang, R. A. Lindquist, J. Moffat, et al. Cellprofiler: image analysis software for identifying and quantifying cell phenotypes. Genome biology, 7(10):1–11, 2006.

Y. Chen, Y. Shi, and B. Zhang. Optimal Control Via Neural Networks: A Convex Approach. In International Conference on Learning Representations (ICLR), 2019.

M. Cuturi. Sinkhorn Distances: Lightspeed Computation of Optimal Transport. In Advances in Neural Information Processing Systems (NeurIPS), volume 26, 2013.

P. Demetci, R. Santorella, B. Sandstede, W. S. Noble, and R. Singh. Gromov–Wasserstein Optimal Transport to Align Single-Cell Multi-Omics Data. BioRxiv, 2020.

A. Dixit, O. Parnas, B. Li, J. Chen, C. P. Fulco, L. Jerby-Arnon, N. D. Marjanovic, D. Dionne, T. Burks, R. Raychowdhury, et al. Perturb-Seq: Dissecting Molecular Circuits with Scalable Single-Cell RNA Profiling of Pooled Genetic Screens. Cell, 167(7):1853–1866, 2016.

C. J. Frangieh, J. C. Melms, P. I. Thakore, K. R. Geiger-Schuller, P. Ho, A. M. Luoma, B. Cleary, L. Jerby-Arnon, S. Malu, M. S. Cuoco, et al. Multimodal pooled perturb-cite-seq screens in patient models define mechanisms of cancer immune evasion. Nature genetics, 53(3):332–341, 2021.

V. A. Green and L. Pelkmans. A systems survey of progressive host-cell reorganization during rotavirus infection. Cell host & microbe, 20(1):107–120, 2016.

A. Gretton, K. M. Borgwardt, M. J. Rasch, B. Schölkopf, and A. Smola. A kernel two-sample test. The Journal of Machine Learning Research, 13(1), 2012.

M. Guizar-Sicairos, S. T. Thurman, and J. R. Fienup. Efficient subpixel image registration algorithms. Optics letters, 33(2):156–158, 2008.

G. Gut, M. D. Herrmann, and L. Pelkmans. Multiplexed protein maps link subcellular organization to cellular states. Science, 361(6401), 2018.

C.-W. Huang, R. T. Q. Chen, C. Tsirigotis, and A. Courville. Convex Potential Flows: Universal Probability Distributions with Optimal Transport and Convex Optimization. In International Conference on Learning Representations (ICLR), 2021.

G.-J. Huizing, G. Peyré, and L. Cantini. Optimal transport improves cell-cell similarity inference in single-cell omics data. bioRxiv, 2021.

L. Kantorovich. On the transfer of masses (in Russian). In Doklady Akademii Nauk, volume 37, 1942.

D. P. Kingma and J. Ba. Adam: A Method for Stochastic Optimization. In International Conference on Learning Representations (ICLR), 2014.

M. Knott and C. S. Smith. On the optimal mapping of distributions. Journal of Optimization Theory and Applications, 43(1), 1984.

A. Korotin, L. Li, A. Genevay, J. Solomon, A. Filippov, and E. Burnaev. Do Neural Optimal Transport Solvers Work? A Continuous Wasserstein-2 Benchmark. arXiv Preprint, 2021.

B. A. Kramer and L. Pelkmans. Cellular state determines the multimodal signaling response of single cells. bioRxiv, 2019.

H. Lavenant, S. Zhang, Y.-H. Kim, and G. Schiebinger. Towards a mathematical theory of trajectory inference. arXiv preprint arXiv:2102.09204, 2021.

P. Liberali, B. Snijder, and L. Pelkmans. A hierarchical map of regulatory genetic interactions in membrane trafficking. Cell, 157(6):1473–1487, 2014.

R. Lopez, J. Regier, M. B. Cole, M. I. Jordan, and N. Yosef. Deep generative modeling for single-cell transcriptomics. Nature methods, 15(12):1053–1058, 2018.

M. Lotfollahi, F. A. Wolf, and F. J. Theis. scGen predicts single-cell perturbation responses. Nature Methods, 16(8), 2019.

A. Makkuva, A. Taghvaei, S. Oh, and J. Lee. Optimal transport mapping via input convex neural networks. In International Conference on Machine Learning (ICML), volume 37, 2020.

L. McInnes, J. Healy, and J. Melville. UMAP: Uniform Manifold Approximation and Projection for Dimension Reduction. arXiv Preprint, 2018.

P. Mokrov, A. Korotin, L. Li, A. Genevay, J. Solomon, and E. Burnaev. Large-Scale Wasserstein Gradient Flows. arXiv Preprint, 2021.

G. Schiebinger, J. Shu, M. Tabaka, B. Cleary, V. Subramanian, A. Solomon, J. Gould, S. Liu, S. Lin, P. Berube, et al. Optimal-Transport Analysis of Single-Cell Gene Expression Identifies Developmental Trajectories in Reprogramming. Cell, 176(4), 2019.

S. M. Shaffer, M. C. Dunagin, S. R. Torborg, E. A. Torre, B. Emert, C. Krepler, M. Beqiri, K. Sproesser, P. A. Brafford, M. Xiao, et al. Rare cell variability and drug-induced reprogramming as a mode of cancer drug resistance. Nature, 546(7658):431–435, 2017.

B. Snijder, R. Sacher, P. Rämö, E.-M. Damm, P. Liberali, and L. Pelkmans. Population context determines cell-to-cell variability in endocytosis and virus infection. Nature, 461(7263):520–523, 2009.

B. Snijder, R. Sacher, P. Rämö, P. Liberali, K. Mench, N. Wolfrum, L. Burleigh, C. C. Scott, M. H. Verheije, J. Mercer, et al. Single-cell analysis of population context advances RNAi screening at multiple levels. Molecular Systems Biology, 8(1):579, 2012.

S. G. Stark, J. Ficek, F. Locatello, X. Bonilla, S. Chevrier, F. Singer, G. Rätsch, and K.-V. Lehmann. Scim: universal single-cell matching with unpaired feature sets. Bioinformatics, 36, 2020.

T. Stoeger, N. Battich, M. D. Herrmann, Y. Yakimovich, and L. Pelkmans. Computer vision for image-based transcriptomics. Methods, 85:44–53, 2015.

A. Taghvaei and A. Jalali. 2-Wasserstein Approximation via Restricted Convex Potentials with Application to Improved Training for GANs. arXiv Preprint, 2019.

S. Van der Walt, J. L. Schönberger, J. Nunez-Iglesias, F. Boulogne, J. D. Warner, N. Yager, E. Gouillart, and T. Yu. scikit-image: image processing in python. PeerJ, 2:e453, 2014.

C. Villani. Topics in Optimal Transportation, volume 58. American Mathematical Soc., 2003.

K. D. Yang, K. Damodaran, S. Venkatachalapathy, A. C. Soylemezoglu, G. Shivashankar, and C. Uhler. Predicting cell lineages using autoencoders and optimal transport. PLoS Computational Biology, 16(4), 2020.

S. Zhang, A. Afanassiev, L. Greenstreet, T. Matsumoto, and G. Schiebinger. Optimal transport analysis reveals trajectories in steady-state systems. bioRxiv, 2021.

